# A Multi-perspective Analysis of Retractions in Life Sciences

**DOI:** 10.1101/2020.04.29.063016

**Authors:** Bhumika Bhatt

## Abstract

The aim of this study is to explore trends in retracted publications in life sciences and biomedical sciences over axes like time, countries, journals and impact factors, and topics. Nearly seven thousand publications, which comprise the entirety of retractions visible through PubMed as of August 2019, were used. This work involved sophisticated data collection and analysis techniques to use data from PubMed, Wikipedia, and WikiData, and study it with respect to the above mentioned axes. Importantly, I employ state-of-the-art analysis and visualization techniques from natural language processing (NLP) to understand the topics in retracted literature. To highlight a few results, the analyses demonstrate an increasing rate of retraction over time and noticeable differences in the publication quality (as measured by journal impact factor) among top countries. Moreover, while molecular biology and cancer dominate retractions, we also see a number of retractions not related to biology. The methods and results of this study can be applied to continuously understand the nature and evolution of retractions in life sciences, thus contributing to the health of this research ecosystem.

## 1 Introduction

The cutthroat competition in academia, the rush to publish or just the greed of career advancement and scientific grants can lead scientists to publish flawed results and conclusions. While some of these errors can be unintentional flaws and honest mistakes, others are intentional scientific misconduct. According to a 2012 report, 67.4% of the then retractions in biomedical research literature were due to misconduct [1]. In this, 43.4% was due to fraud or suspected fraud,14.3% due to duplicate publication, 9.8% due to plagiarism and rest due to unknown or other reasons. Another 2018 study found that the most common reasons for retraction in open access journals were errors, plagiarism, duplicate publication, fraud/suspected fraud and faked peer review process [2]. Flawed publications not only undermine the integrity of science but can also lead to the propagation of erroneous data to other genuine publications, and thus retraction of such work is a necessary step to preserve the integrity of the entire scientific research ecosystem. There exist a few previous publications to understand the reasons, scope, and impact of retractions in life sciences [3, 4, 5, 6, 7, 8, 9, 10, 11, 12, 13, 14, 15, 16, 17, 18]. Researchers have analyzed retractions by intentions (i.e., mistakes, fraud, fabrication, etc.) as well as studied trends over time, countries, journals and in relation to authors. Much of this work has been manual and cannot benefit significantly from available computational techniques. My work in this manuscript differs from the previous works in both the end goals and the approach. I map the available retraction data to various dimensions such as countries, journals and impact factors, time and subject areas and discover previously unknown or unconfirmed trends in the data. Studying reasons behind retractions was out of scope of this work. This study has addressed several challenges, most of which stem from the unstructured nature of the data. The affiliations and abstracts for publications are all unstructured. There is no easy way to obtain the authors’ countries, for instance, which we need in our analysis. Consequently, I had to gather country data from other sources and join it with the PubMed data using non-trivial automatic and manual analysis. I also perform topic modeling on the abstracts to understand and characterize retracted literature by subject areas. This involved natural language processing, analyses and visualization. Finally, the entire work has involved significant data cleaning and processing to convert the data into a form amenable for analysis and drawing conclusions. In the next few sections, I discuss the methodology along with the findings in greater detail. This work makes the following major contributions.

- *Large dataset.* This manuscript reports on all retractions until August 2019, as reported on PubMed. To the best of the author’s knowledge the present study has used the biggest dataset among all retraction studies so far.
- *Trends by time and journal impact factors.* The work shows that retraction rate is increasing over time and retraction differs significantly by journals, with most retractions happening in lower impact factors.
- *Trends by countries.* I show that authors from India and China have more of their retractions in lower impact factor journals while authors from Germany, Japan, and the US have more of their retractions in higher impact factor journals when compared to India and China.
- *Trends by subject areas.* This work implements a topic model based on statistical techniques to identify the subject areas of retracted publications. While molecular biology, containing genetics and biochemistry, and cancer dominate the topics among retracted papers, we also find a significant number of papers that are not related to life sciences.

## 2 Methods

I collected the data for retracted publications from PubMed on August 26, 2019 using the Eutilities API [19]. From this, I extracted articles’ titles, journal names, publication and retraction dates, authors’ affiliations, and abstracts. The extraction and processing was done in Python. The final dataset contains 6,936 unique retracted articles. Another possible source for retraction data is the Retraction Watch database [20], which is a curated dataset. Between these two sources, I used PubMed as it also contain article abstracts, which I use for this study.

### 2.1 Processing Dates

The publication dates can be found most often as PubDate (a field in Eutilities XML) but may also be available as MedlineDate or PubmedPubDate fields. I chose the minimum of these dates. The 832 entries with the day of month missing were assigned 15 (mean number of days in a month) as the day. Another 58 entries, which were missing the month, were dropped for the time-based analysis. Retraction dates are available under the CommentsCorrections field as unstructured text, which was parsed using regular expressions. Most (4,211) entries were missing the day, which was again assigned with 15. 709 entries had imprecise or missing retraction month or year and were dropped. In 23 cases the retraction date was earlier than the publication date. Further investigation revealed that the correct publication date was not available for them. All such entries were also dropped. Additionally, in seven other cases, the retraction date appeared to be before the publication date but this was simply due to artificially assigning the day to 15. Note that entries dropped here are still used for non-time-based analysis.

### 2.2 Processing Country Names

Author affiliations can be used to identify which countries the authors belong to. However, this poses a few challenges.

- Affiliations are unstructured and do not follow a common format, it is inherently challenging to derive structured information such as countries from them.
- A country may be known by multiple names, for example, Germany and Deutschland; and Brazil and Brasil. These examples appear in our dataset.
- A country name may not be present in an affiliation but instead must be inferred. For example, if an affiliation mentions “Stanford University” only, it must be inferred that the country is United States.

I addressed the first challenge through an algorithm that scans an affiliation in the reverse order. It uses heuristics to tokenize it into groups of words, which it then uses to identify or infer a country. To solve the second challenge, I obtained a list of alternative country names from Wikipedia. When looking for countries, I searched for all country names, including the alternative ones. The third challenge is more difficult. It would be desirable to have a mapping of all the universities and organizations in the world to their home countries. Fortunately, an approximation of this mapping is possible through ontological databases such as WikiData, which contain crowd-sourced information relating various entities and can be queried to obtain desired mappings. Using WikiData’s SPARQL (a query language) interface, I obtained a list of 84,476 organizations including universities, research institutes, engineering colleges, university systems, international organization, hospitals, businesses, research centers, and academies of sciences and their corresponding countries. While certainly not exhaustive, it would likely cover most of the affiliations we are likely to see in our dataset. Moreover, while neither the list of alternative country names nor the list of organizations may be exhaustive, together they would provide better resolution of countries than those provided by one of these methods alone. After processing affiliations in such an automatic fashion, I manually checked them (about 100 were updated). My methodology uses all countries appearing in affiliations, counting each country at most once for a given paper. 25 cases had no individual authors but only collective groups as authors. I manually identified the countries these groups belong to. The data also lacked affiliations for 439 retracted papers.

### 2.3 Journals

The journal names from the retraction data were combined with impact factors, which were obtained from 2019 edition of Journal Citation Reports (JCR) [21]. Before matching the journal names to those in the impact factor list, the names were normalized by making them lowercase, removing spaces and punctuations, and so on. The matching is still not trivial: for instance, an original journal name in the PubMed data is ‘JAMA’, which is available in the impact factor list as ‘JAMA-Journal of the American Medical Association’. Such mismatches were identified and fixed manually.

Furthermore, a few journals, e.g., Biochimica et Biophysica Acta, have several sections, each of which is separately mentioned in JCR. However, the PubMed lists the journal only without the section. In these cases, I manually resolved the sections to assign the right impact. The two manual steps allowed assigning impact factor to 153 additional articles.

### 2.4 Topic Modeling

Understanding the subject areas and themes of the retracted articles is a major effort in this work. An obvious approach is to use MeSH [22] identifiers, which denote both broad subject areas and detailed topics and are manually curated for each article. Their primary purpose is to enable efficient search for articles [23, 24]. One may suggest that we could just use the MeSH to identify the themes in articles. Unfortunately, this does not work for us for at least two reasons: (a) there are 1,713 articles (i.e., about a quarter of the dataset), which do not have any qualifiers assigned; and (b) MeSH provides multiple qualifiers for a given article but does not indicate their relative importance, thus making it difficult to identify a single, most-important subject area for the article. An alternative is performing topic modeling, which uses statistical techniques to discover abstract “topics” in given documents (here, retracted literature).

Topic models learn topics from a corpus in an unsupervised way. Latent Dirichlet Allocation (LDA) [25]] is a state-of-the-art method used for topic modeling. It is a generative probabilistic topic model, based on the idea that each document is a probability distribution over topics and each topic is a distribution over words. Thus, hidden topics in documents can be found out from the collection of words that co-occur frequently. LDA has been used in a number of natural language processing and information retrieval tasks and has even been explored in life sciences literature. For instance, it was used to discover relationships among drugs, genes, proteins, and pathways in several articles [26, 27, 28, 29]. This work, however, uses LDA to understand the broad topics appearing in the literature under consideration. Next, I discuss data preprocessing to obtain the “words” for the model, and building the LDA model to understand the topics.

#### 2.4.1 Data Preprocessing

This step was used to clean and augment the available data to allow building an effective model. Articles that either lacked an abstract or had an abstract describing only their retraction were excluded, resulting in 6,417 articles. Wherever possible, abbreviations were unabbreviated. I used a simple heuristic for this – upon encountering a possible abbreviation within parentheses, its letters are matched with the first letters of the immediately preceding words. If a match is found, this abbreviation is expanded all over the abstract. This heuristic only helps with the common case and may miss cases like ‘hookworm’ abbreviated as ‘HW’ and ‘tuberculosis’ abbreviated as TB.

##### Tokenization and lemmatization

This step broke text into words, removed some words based on their part-of-speech usage, and then normalized. Part of speech (PoS) tagging is used to enable lemmatization or normalization of words. For this work, part of speech tagging was additionally used to extract only nouns, proper nouns, and adjectives (e.g, ‘cell’, ‘Parkinson’s disease’, and ‘pulmonary’), while ignoring other parts of speech, such as verbs (e.g., ‘examined’ and ‘discovered’), which do not appear useful to our current task. The obtained words were then lemmatized to extract roots of the words. This normalizes the words according to their use as different parts of speech. For instance, the plural ‘symptoms’ would be transformed to the singular ‘symptom’. As life science literature frequently contains phrases, such as ‘cell line’, ‘reactive oxygen species’, which have domain-specific connotations not conveyed by the comprising words such as ‘cell’ and ‘line’ (when considering ‘cell line’), I augmented the derived terms with such phrases. To identify these phrases, I used Named Entity Recognition. The techniques discussed here were implemented through Scispacy [30], a python library extending spaCy [31] for processing scientific and biomedical texts.

##### Stopword elimination

Stopwords are words that do not have enough meaning to differentiate between two texts. The NLTK [32] library contains 179 stopwords. I further added more words such as ‘proof’, ‘researcher’, ‘record’ that have no differentiating effect in life sciences literature, making the stopword list 544 words long. Moreover, I made all words in lowercase to make the subsequent steps case-insensitive and removed all words consisting of digits only or one letter only. In addition, symbols that are common in scientific literature such as =,<,>,− were removed.

#### 2.4.2 Building an LDA model

I now discuss the implementation of the LDA model. The first step is to construct a vocabulary from the terms derived from data processing. Only those terms that appeared in at least twenty articles and in not more than 15% of the abstracts were used. The rationale is that terms appearing in too few or too many documents would not convey any meaningful pattern. This vocabulary is used to construct a document-term matrix, where each row corresponds to an article abstract (documents) and each column corresponds to a word or term. Each cell in the matrix represents the number of occurrences of the term in the document. This is also known as a bag-of-words (BoW) model and is an input to the LDA algorithm. LDA provides as many topics K as defined by the user. A low K can provide broad topics while a high K can give topics with words repeated in multiple topics, thus making them difficult to interpret. To arrive at an appropriate K, I started with a target K = 15 and built models for values of K around it to identify the model with the most interpretable topics. I finally selected K = 16. This implementation was done using the Gensim library [33]. The model was visualized with LDAvis [34, 35].

## 3 Results

### 3.1 Trend over time

As seen from the red curve in Figure 1a, the number of retractions for the newly published papers has been increasing each year. To answer whether this rise can be attributed to the increasing rate of publications, I collected per-year publication statistics from PubMed and plot the retraction rate in Figure 1b. The retraction rate per 10,000 publications was 0.38 in 1985 (here, a retraction is attributed to the year of publication) and it rose to 2.03 in 2000 and 5.95 in 2014. Next, I checked how many publications are retracted in a given year irrespective of their publication year. From the blue curve in Figure 1a, it can be confirmed that starting mid to late 2000s there is a steady increase in number of retractions. Figure 1c plots the time to retraction. Maximum retractions happen within 1 year of publication and the number decreases as years pass by. It takes, on an average, 3.8 years for a publication to be retracted (with standard deviation of 4.01 years) and this explains why we see less number of retractions for the papers published in year 2015 onwards compared to the year 2014 (Figure 1a). Additionally, the 25th percentile for years to retraction is about 1 year and the 75th percentile is 5.3 years. The median is 2.3 years.

**Figure 1:**
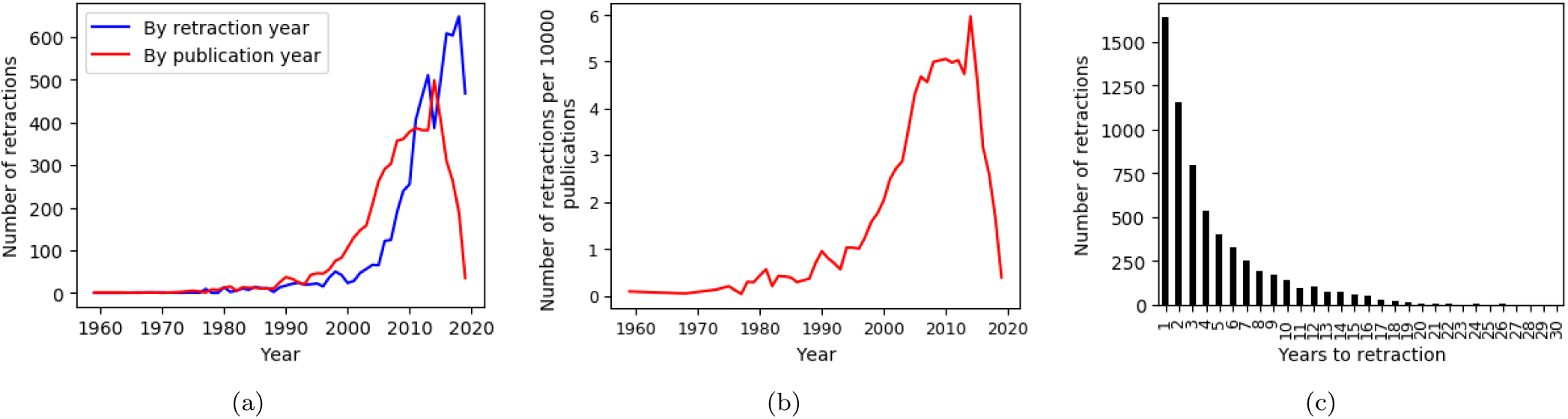
Retraction trend over time. (*a*) The number of retractions show a generally increasing trend both by the year of publication (red curve) and by the year of retraction (blue curve). Note that the red curve falls off in recent years because there will likely be more retractions in future among publications published in recent years. The blue curve falls off in 2019 because we have only partial data for 2019. (*b*) Similar analysis as (*a*), except the number retractions is per 10,000 publications. This shows that even after accounting for the increasing number of publications, the rate of retraction is increasing. (*c*) Most retractions happen soon after publication.

### 3.2 Trend among countries

In the available data, the authors come from 98 countries. United States, China, Japan, India, and Germany occupy the first five ranks (Figure 2a). Focusing more on these 5 countries revealed that retracted publications from China soared in mid 2010’s especially for the years 2014-2015 (Figure 2b). This increase in retractions correlates well (pearson correlation coefficient = 0.9 for 2000-2015) with the increase in publications from China (Figure 2c). Next, I considered countries by their retraction rate. In order to avoid getting countries that do not publish actively, I set a threshold that a country should have at least 10,000 publications. The countries with highest retraction rates are Iran, Tunisia, Pakistan, Bangladesh and India (Figure 2d). I also compared the distribution of retracted literature over the journal impact factor (IF) for the top five countries. For lower IF retractions, China dominates while at higher IFs, the USA dominates (Figure 3a). Figure 3b makes a deeper analysis. India’s retractions in journals with no IF are at 33% of its overall retractions, which is highest percentage among these five countries. Similarly, the highest percentage for retractions in journals with IF 0-5 is for China and for journals above 5 is for the USA. Additionally, Japan and Germany also have lower percentage of their retractions in low IF journals (no impact factor and journals with 0–5 impact factor) compared to India and China. This shows a slight skew of distribution of retracted papers for these three developed countries (USA, Japan, Germany) showing more of their retractions in higher impact factor journals compared to the two developing countries, China and India.

**Figure 2:**
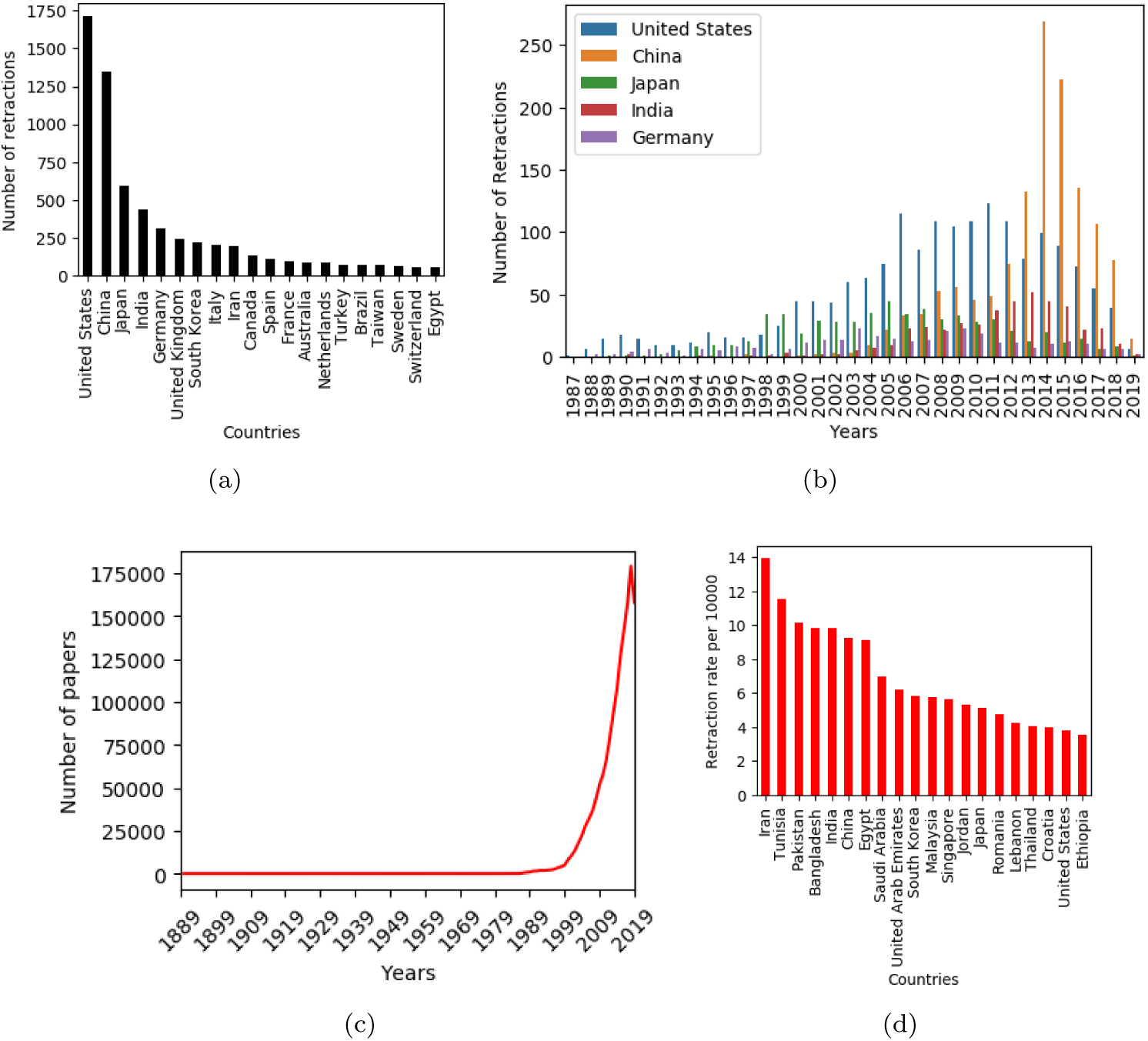
Retraction trend among countries. (*a*) Top 20 countries with highest number of retractions. (*b*) Retraction trend over time for the five countries with highest retractions. (*c*) Publication over years for China showing very high increases in recent decades. This is a factor for the high number of retractions for China in recent years. (*d*) Top 20 countries with highest retraction rate per 10,000 publications.

**Figure 3:**
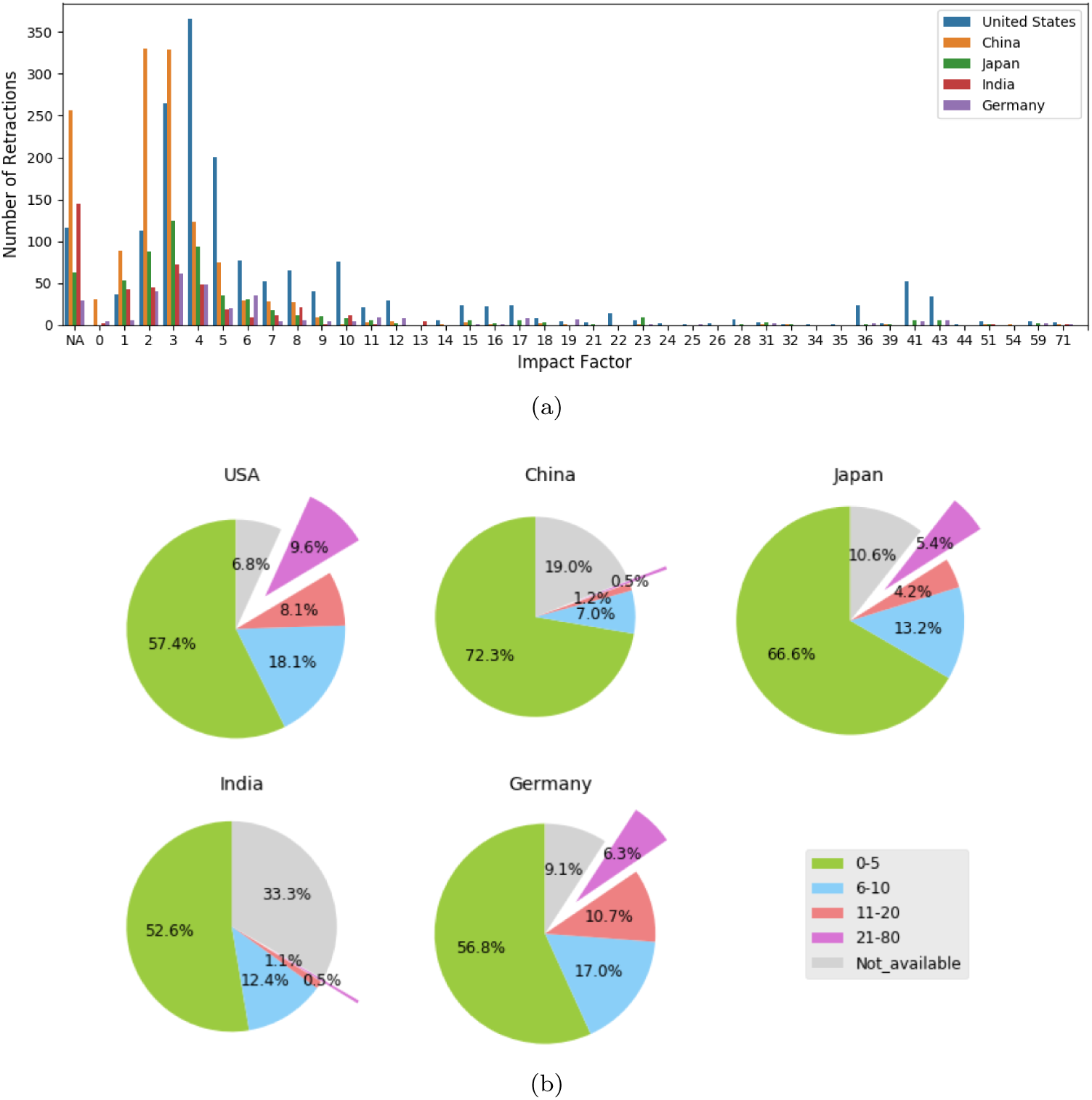
Distribution of retracted publications over their journal’s impact factor for the top five countries with highest retractions. (*a*) Number of retracted publications by their corresponding journal’s impact factor. Note: NA impact factor indicates journals without impact factor. (*b*) Pie charts indicating individual country’s distribution of retracted publications according to impact factor. Impact factors have been clustered into five groups: Not available, 0-5, 6-10, 11-20, 21-80.

### 3.3 Trend among journals

The 6,936 papers analyzed here were published in 2,102 different journals. 54% of these journals have only one retracted paper (Figure 4a). The highest retractions are from Journal of biological chemistry with 279 retractions, Plos One with 177 and Tumor Biology with 145 retractions (Figure 4b). In addition, out of the top 21 journals with the highest retractions (the 20th journal is tied with the 21st), Diagnostic pathology (total 39 retractions) followed by Tumor Biology has the highest retraction rate (number of retractions per total number of papers published by a journal) among these 21 journals with highest retractions(Figure 4c). Next, (Figure 4d) analyzes retractions by journal impact factor (IF), rounded to the nearest integer. With 1,273 retractions, journals with factor of 3 had the most retractions.

**Figure 4:**
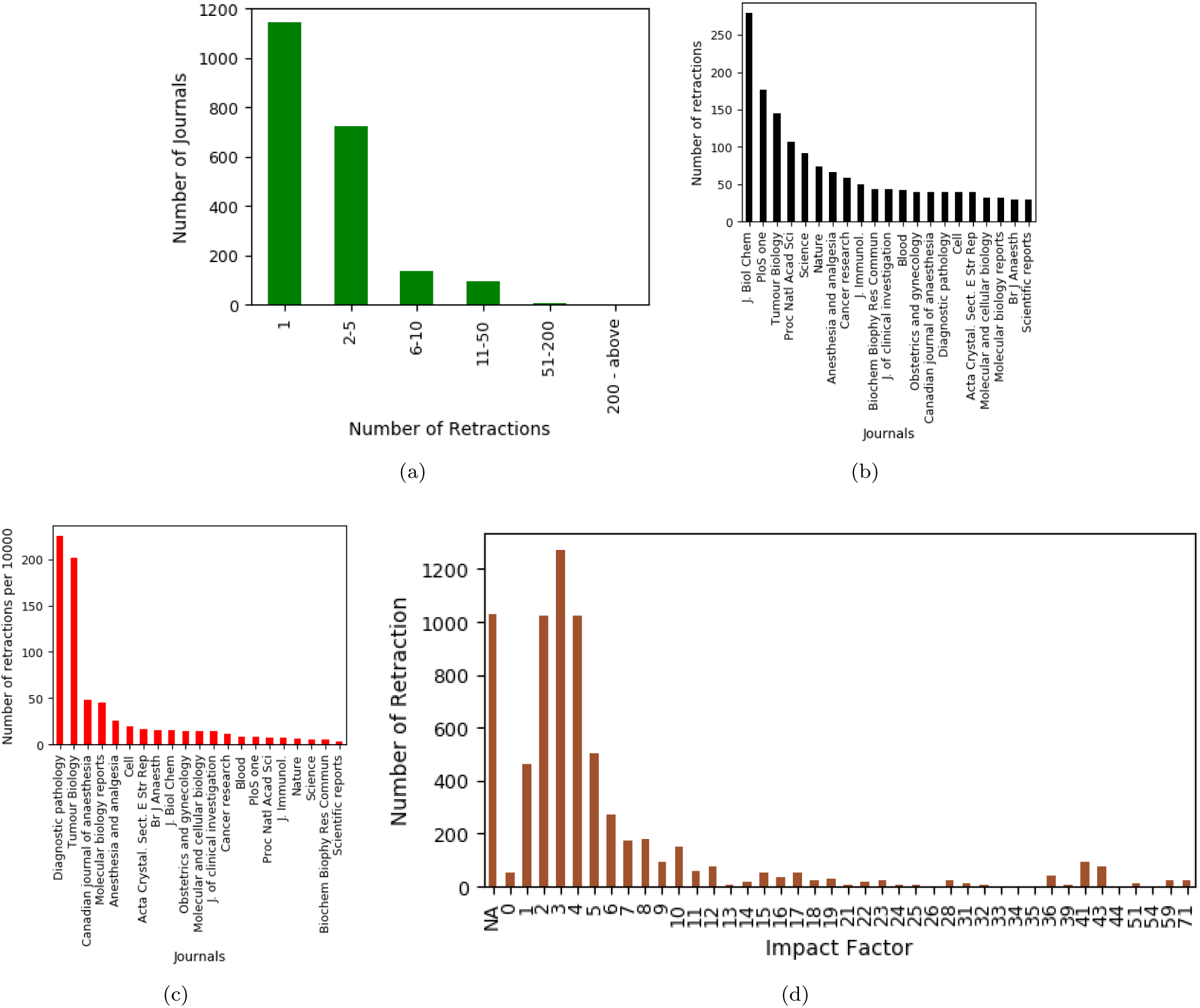
Retraction trend among journals. (*a*) Number of journals against the number of papers retracted from them. (*b*) Top 21 journals with highest number of retracted publications. (*c*) Retraction rate per 10,000 publications for journals in (*b*). (*d*) Number of retractions coming from journals with different impact factors (NA - Not available impact factor).

This was followed by journals that had no impact factor, which is closely followed by papers with IF 2 and IF 4. 55% of the retractions are from journals with IF below 3.

### 3.4 Topic analysis

#### Visualization of results

Figure 5 presents a screenshot of the interactive visualization prepared with LDAvis of the LDA model with 16 topics. Topics are represented as circles. Closely related topics are spatially close and conversely, unrelated topics are distant. Additionally, bigger circles represent more prevalent topics.

**Figure 5:**
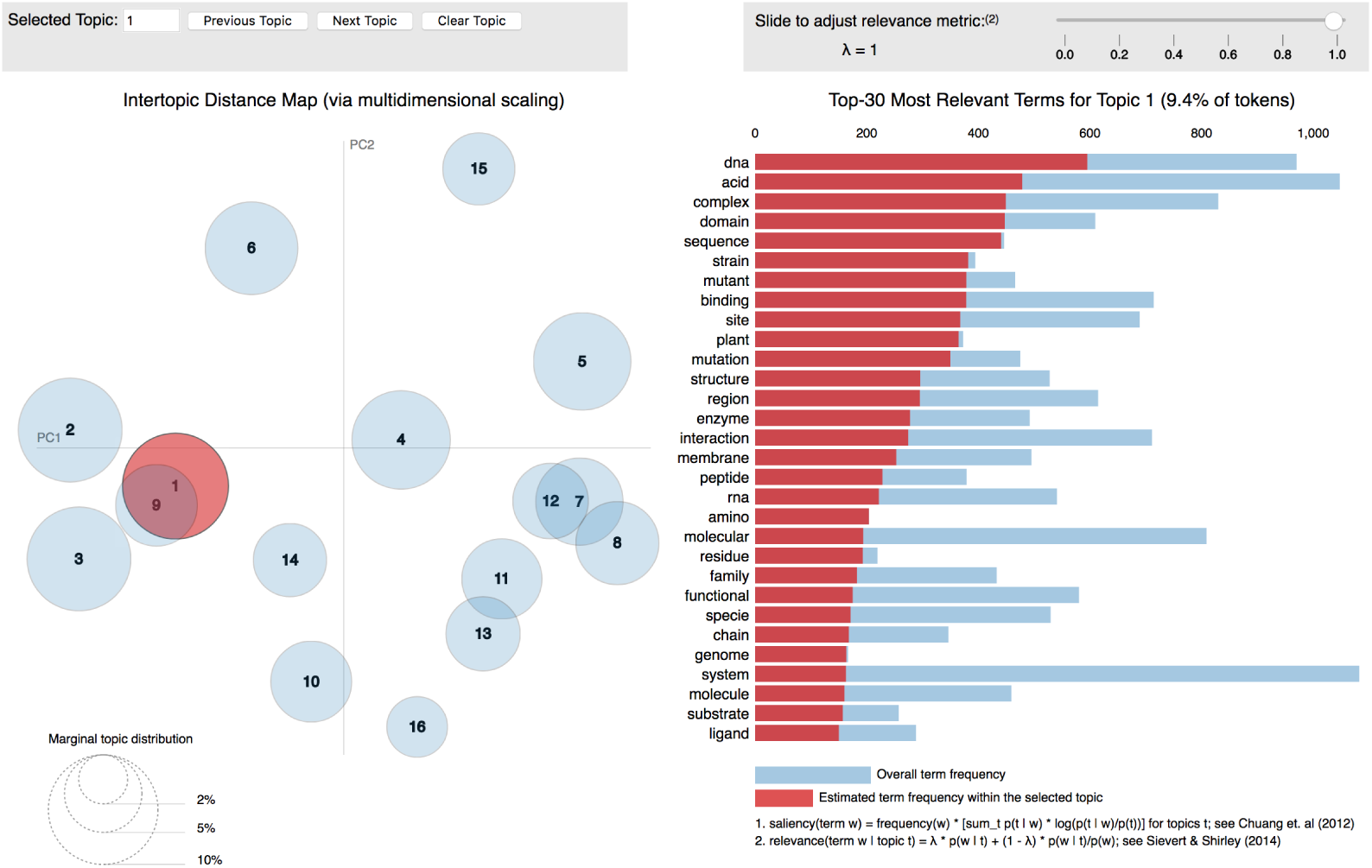
Visualization of topics for an LDA model with 16 topics.

#### Topic interpretation

LDA provides topics in the form of co-occurring words. These topics need further interpretation. I analyzed these topics based on the words occurring within them as well as the abstracts that had high prevalence of the respective topics. Table 1 presents the topic analysis: the topic interpretation, the salient keywords representing the topic and remarks indicating how the abstracts are relevant to the interpreted topic.

**Table 1:**
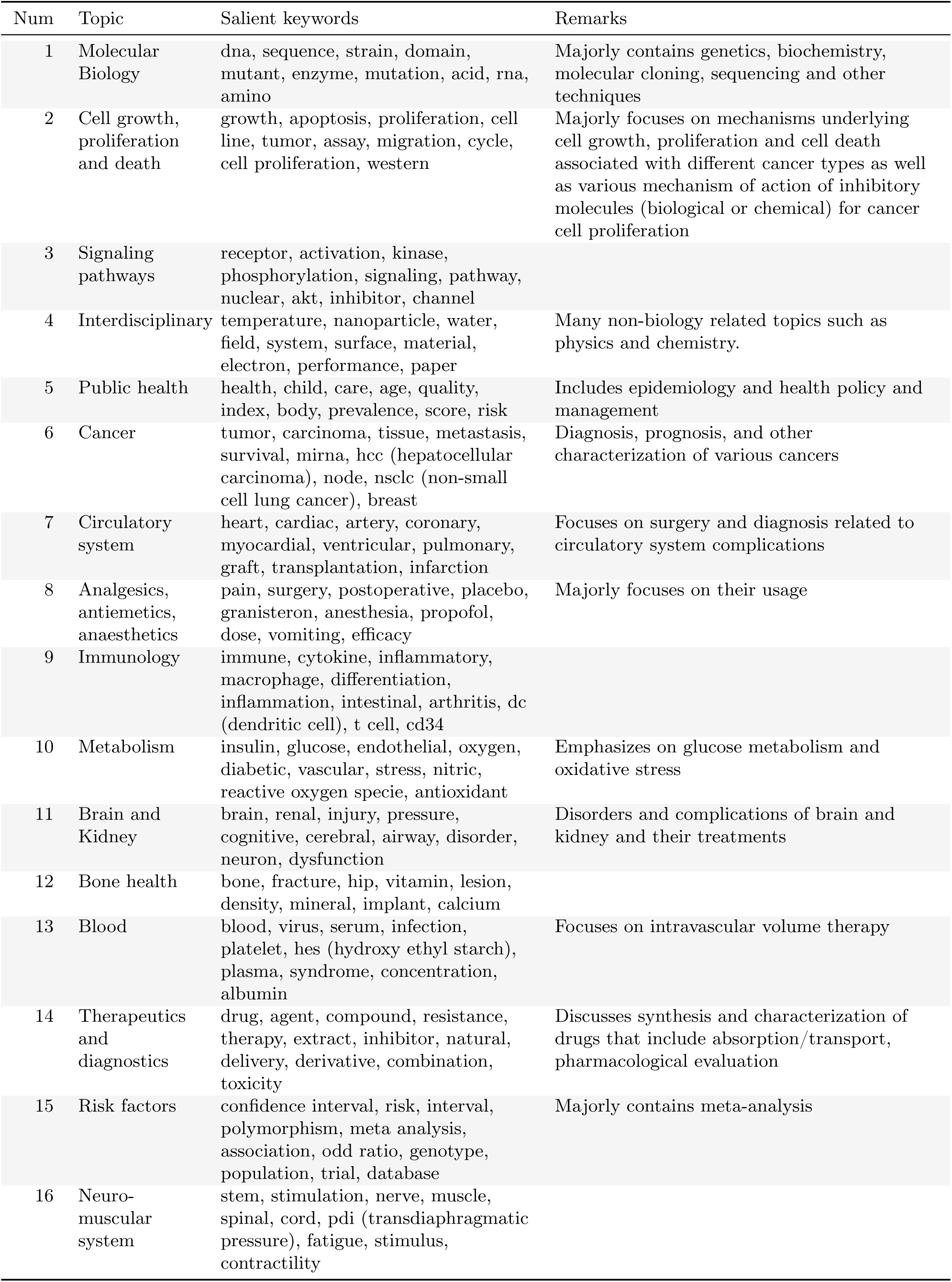
Topics for the constructed LDA model.

#### Analysis

Note that some topics are more closely related than others. For example topic 1 and topic 9, which are about molecular biology and immunology respectively, show an overlap in Figure 5. The topics have some common terms such as complex, region, peptide, interaction, and membrane. Also, molecular biology is a broad topic encompassing biomolecules that are functional in various aspects of cell biology, including in immunology, making the overlap intuitive. Similarly, we find topic 1 closely related to signaling pathways (topic 3) and cell growth, proliferation and death (topic 2) and therefore, these topics appear close to each other in Figure 5. For another example, topic 15 (risk factors), which frequently concerns genetic epidemiology, is related to public health (topic 5), which contains epidemiology, and the two topics are not far from each other. Additionally, topic 7 (circulatory system) overlaps with topic 12 (bone health) as well as with topic 8 (analgesics, anti-emetics, anaesthetics), all related to human body and health. Similarly, we observe an overlap between topic 11 (brain and kidney) and topic 13 (blood). Recall that a single document (abstract) can be made up of multiple topics, but the prevalence (represented as a probability) of these topics in a document can vary. This leads to two ways of ranking the topics:

- By number of documents where the given topic is the most prevalent (Figure 6a).
- By the total prevalence across all documents (Figure 6b).

**Figure 6:**
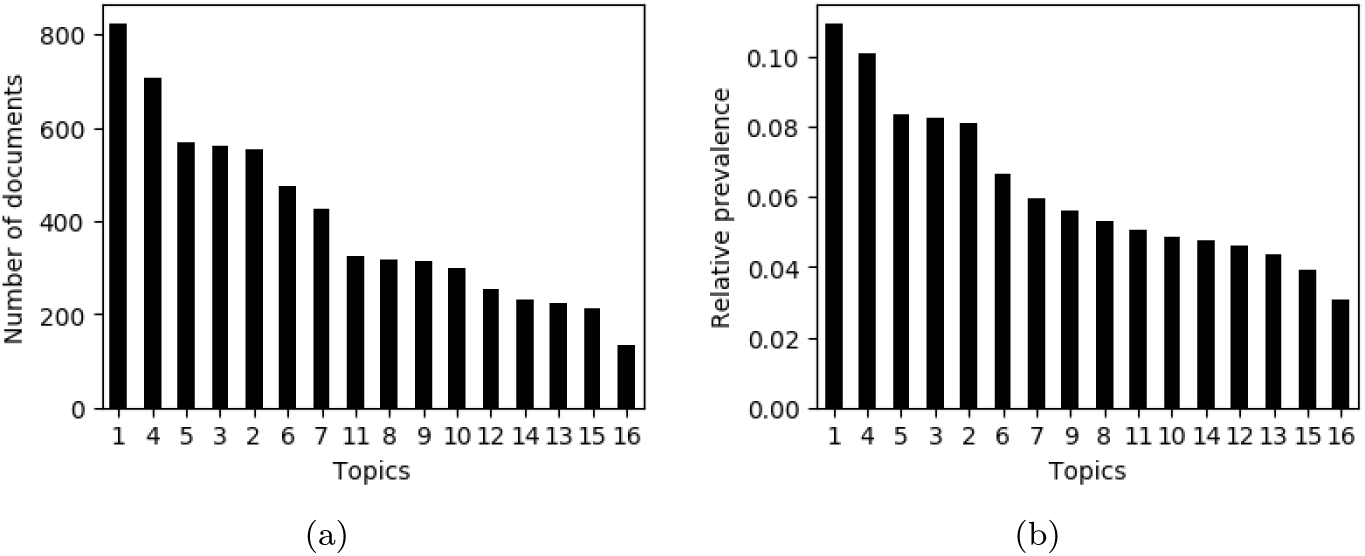
Ranking of topics according to (*a*) the number of documents where they have highest probability compared to the other topics; and (*b*) relative prevalence.

Topic 1 (Molecular biology) is the top topic according to both the ranking criteria. This is intuitive because the topic is quite vast, encompassing genetics and biochemistry. Topic 4 ranks second by both rankings. Further study reveals that while this topic contains interdisciplinary articles such as on biophysics and computational biology, many of the articles are not related to biology at all. Upon checking 100 randomly selected abstracts of the 705 comprising this topic, I found 49 to be non-biology. Extrapolating this to the rest of 705 abstracts, this suggests that about 5% (345 out of 6417) abstracts can be non-biology related. Interestingly, topic 2 (cell growth, proliferation and death) is highly related to cancer. Among randomly selected 100 abstracts from 555 abstracts that have topic 2 as their most prevalent topic, 77 were related to cancer. If we extrapolate this percentage of abstracts related to cancer, then 427 abstracts would be cancer-centric. Adding this number to the number of abstracts from cancer (topic 6), i.e., 474, the total abstracts that are related to cancer would be 901. This would make cancer the most prevalent topic in all the documents.

## 4 Discussion

The increase in retraction rate in Figure 1b implies an increasing number of retractions even after accounting for the increase in the number of publications. Previous studies have also reported such findings [6, 7, 8]. Note that the present study uses the largest dataset for such analysis and still reports this finding consistent with previous studies.

Although about six retractions over 10,000 publications (Section 3.1) may appear small, the disruptive potential of papers, many claiming breakthroughs, including in clinical trials and medical treatments, cannot be underestimated. This trend may be due to increased fraud, mistakes, and so on but may also be due to increased vigilance.

While it is important to understand retraction trend among countries based on their absolute number of retractions, with differences in their research publication productivity it becomes crucial to understand retraction rate as well. In our analysis, Iran has the highest retraction rate. Masoomi and Amanollahi called out scientific fraud in Iran, citing scientific misconduct behind 80% of Iran’s retractions [36].

We observed impact factor distribution differences among the countries with the highest retractions. This empirically corroborates Fang et al. [1], that fraud is more prevalent in countries with long-standing research culture, e.g., U.S., Germany, Japan, and among high-impact journals while plagiarism and duplication is often present in countries still developing such culture and is often associated with low-impact journals.

In our topic model, some topics appear very broad and some noise is creeping in the topics. Noise in LDA models is not unheard of and is especially an artifact of small data like in this study, which comprised only abstracts from only about six thousand articles. Syed and Spruit suggest that using full text (which this study did not have access to) instead of abstracts will largely alleviate the above-mentioned problems of noise and broad topics [37]. Notwithstanding, our model also revealed highly interpretable topics.

In our topic analysis, we see non-biology articles in our dataset. This is because the journals where these articles are published are either broad scientific journals (such as Plos One, PNAS, and Nature) or generic chemistry or physics or computer science (such as The Journal of Chemical Physics, Langmuir, Neural Networks) that sometimes publish articles relevant to life sciences and biomedical sciences. In either case, PubMed indexes the entire journals irrespective of the article subject areas.

Although I have studied variation of retractions with impact factor and with topics, I have not studied how retraction rate varies with these factors. Studying retraction rate would require classifying all published research (or a proxy) by impact factor and topics. This is a daunting task and design of such a study is left for future work.

The missing data may present a threat to validity of the results. A small amount of data – 11% for time-based analysis, 6% for country analysis, and 7% for topic modeling – is missing. Nonetheless, assuming that the data is missing randomly, I believe that the results are statistically valid and generalize to the real situation.

Increased digitalization has obviously led to increased vigilance, e.g., through automatic plagiarism checks and easier access and searchability to articles. Consequently, papers that would have gone without raised eye brows in the past have greater chance of being retracted today. At the same time, the “publish or perish” adage is truer now than ever before. I believe it is therefore important to continuously track and analyze retractions in this evolving landscape and apply the derived insights to improve the integrity of scientific research.

## Author Bio

Bhumika Bhatt is an independent researcher interested in biology and data science. She holds a Ph.D. in biochemistry and molecular biology from Medical University of Vienna, Austria. Her Ph.D. work at Max F. Perutz Laboratories helped develop a new method for detecting short-lived protein-protein interactions in yeast. She was also a postdoctoral fellow at Sanford Burnham Prebys Medical Discovery Institute in San Diego. She runs a blog about biology and data science at https://bhumikabhatt.org.

